# Biased competition in semantic representation during natural visual search

**DOI:** 10.1101/658096

**Authors:** Mohammad Shahdloo, Emin Çelik, Tolga Çukur

## Abstract

Humans divide their attention among multiple visual targets in daily life, and visual search gets more difficult as the number of targets increases. The biased competition hypothesis (BC) has been put forth as an explanation for this phenomenon. BC suggests that brain responses during divided attention are a weighted linear combination of the responses during search for each target individually. Furthermore, this combination is biased by the intrinsic selectivity of cortical regions. Yet, it is unknown whether attentional modulations of semantic representations of cluttered and dynamic natural scenes are consistent with this hypothesis. Here, we investigated whether BC accounts for semantic representation during natural category-based visual search. Human subjects viewed natural movies, and their whole-brain BOLD responses were recorded while they attended to “humans”, “vehicles” (i.e. single-target attention tasks), or “both humans and vehicles” (i.e. divided attention) in separate runs. We computed a voxelwise linearity index to assess whether semantic representation during divided attention can be modeled as a weighted combination of representations during the two single-target attention tasks. We then examined the bias in weights of this linear combination across cortical ROIs. We find that semantic representations during divided attention are linear to a substantial degree, and that they are biased toward the preferred target in category-selective areas across ventral temporal cortex. Taken together, these results suggest that the biased competition hypothesis is a compelling account for attentional modulations of semantic representation across cortex.

**Significance Statement**

Natural vision is a complex task that involves splitting attention between multiple search targets. According to the biased competition hypothesis (BC), limited representational capacity of the cortex inevitably leads to a competition among representation of these targets and the competition is biased by intrinsic selectivity of cortical areas. Here we examined BC for semantic representation of hundreds of object and action categories in natural movies. We observed that: 1) semantic representation during simultaneous attention to two object categories is a weighted linear combination of representations during attention to each of them alone, and 2) the linear combination is biased toward semantic representation of the preferred object category in strongly category-selective areas. These findings suggest BC as a compelling account for attentional modulations of semantic representation across cortex in natural vision.

## Introduction

In daily life, humans frequently search for a multitude of objects in their visual environment. Yet, attending to multiple objects becomes more difficult as the number of targets increases. Psychophysical studies showed that reaction time and error rate during visual search systematically increase with growing number of items to be attended (Eckstein et al., 2000; Wolfe, 2012; Reynolds and Chelazzi, 2004; Luck et al., 1997). The biased competition hypothesis (BC) has been proposed to account for this performance decline (Duncan, 1984). BC reasons that the brain has limited representational capacity. Thus, simultaneous search for multiple visual objects results in a competition among their representations. Moreover, task demands due to spatial (Keitel et al., 2013; Kastner et al., 1998), feature-based (McMains and Kastner, 2011; Bichot et al., 2005; Boynton, 2005), and object-based (Gentile and Jansma, 2010; Reddy et al., 2009) attention can bias this competition in favor of the target (Desimone, 1998).

Several neuroimaging studies provided evidence for competition among cortical representations of multiple objects across visual cortex in the absence of specific task demands (Kastner et al., 1998; MacEvoy and Epstein, 2009; Gentile and Jansma, 2010; Nagy et al., 2011; Baeck et al., 2013; Jeong and Xu, 2017). Gentile and Jansma (2010) measured average BOLD responses in fusiform face area (FFA) while subjects viewed a single face image or a pair of face images. Response to a pair of faces was lower than the summation of the responses when each of the faces were presented individually. Similarly, Nagy et al. (2011) presented four equispaced isolated images of faces or noise images. The number of face images was increased systematically from zero to four. They reported that responses in FFA and lateral occipital complex (LOC) to multiple faces were lower than the summation of the responses to individual faces. MacEvoy and Epstein (2009) suggested a linear model of representational competition among objects. The authors presented either a single or a pair of isolated images of objects from four categories (shoes, chairs, cars, or brushes). Using multivoxel pattern analysis, they showed that the response pattern in object-selective areas in ventral temporal cortex when subjects viewed pairs of objects can be approximated by the mean of response patterns when they viewed each of the objects in isolation. These results were interpreted to imply the existence of competition among representations.

Recent studies also provided evidence for top-down influences in BC during divided attention to multiple objects (Reddy et al., 2009; Gentile and Jansma, 2010). Reddy et al. (2009) studied BOLD responses while subjects attended to a single or a pair of object categories among four alternatives (faces, houses, shoes, or cars). A multivoxel pattern analysis in category-selective areas in ventral temporal cortex revealed that the response pattern during divided attention to two object categories was a weighted linear combination of response patterns while attending to individual targets. Furthermore, the authors reported that in PPA (preferentially responsive to houses) and in FFA (preferentially responsive to faces), combination weights were biased toward the preferred object category. Similarly, Gentile and Jansma (2010) reported that during attention to one of the faces in a pair of face images, FFA responses were biased toward the responses recorded when the target was presented in isolation.

Previous studies on BC have provided evidence for competition in representation of isolated static objects. Yet, natural scenes are intrinsically dynamic and cluttered with many objects and actions. It has recently been suggested that thousands of visual categories are embedded in a continuous semantic space across cortex (Huth et al., 2012), and that category-based visual search causes broad modulations in these semantic representations (Çukur et al., 2013). It is currently unknown whether BC can account for modulations in semantic representation during natural visual search for object categories.

Here we questioned whether BC can account for semantic representation of object-action categories during divided attention in natural visual search. To address this question, we conducted a functional magnetic resonance imaging (fMRI) experiment (Fig.1). Five human subjects viewed 72 min of natural movies while performing three separate tasks in different runs: attend to “humans”, attend to “vehicles”, and attend to “humans and vehicles” (i.e., divided attention). Whole-brain BOLD responses were recorded and category responses for 831 objects and actions were measured separately for each task and each individual subject (Nishimoto et al., 2011). Semantic tuning was estimated via principle component analysis on the response profiles. To test whether the semantic tuning during divided attention can be approximated by a weighted linear combination of the tuning during single-target tasks, ordinary least squares was used among semantic tuning profiles for the three tasks. To reveal the interactions between the attentional bias in semantic tuning and the intrinsic selectivity of brain areas, the semantic tuning distribution during divided attention was regressed onto tuning distributions during the two single-target tasks.

**Figuer 1.**
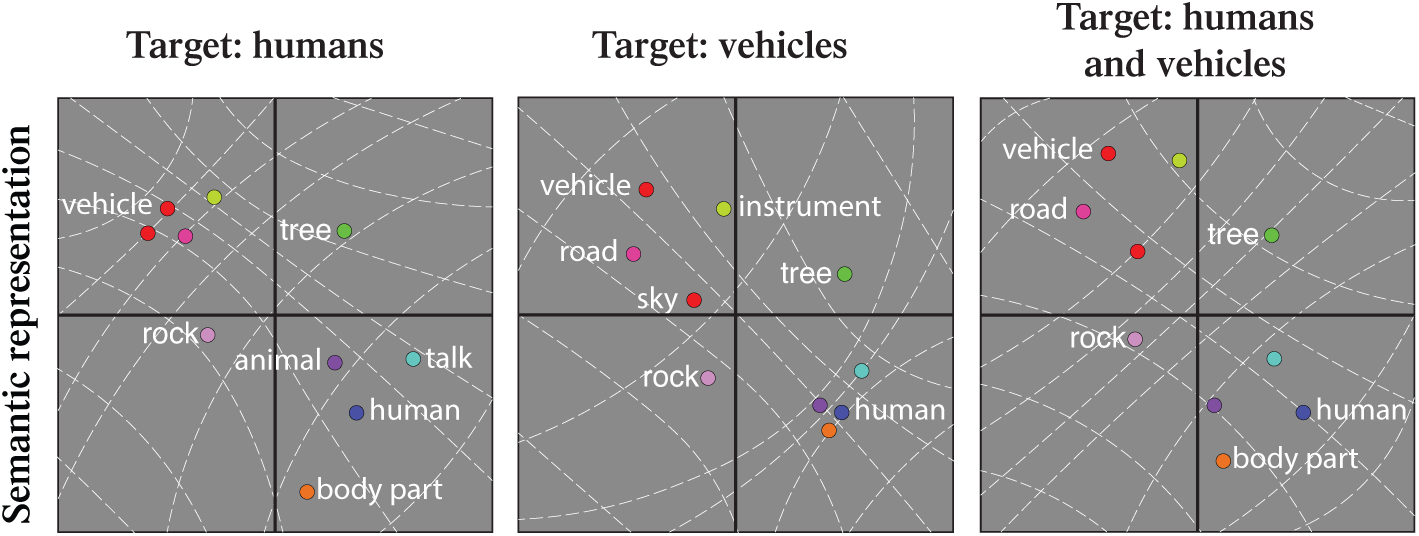
Hypothesized changes in semantic representation during divided attention. Previous studies have proposed that the human brain represents thousands of object and action categories by embedding them in a low-dimensional space based on their semantic similarity. It has been shown that attention warps semantic representation in favor of the target and semantically similar nontargets. Biased competition hypothesis predicts that semantic representation during divided attention is a weighted linear combination of representations during attention to individual targets.

## Materials and Methods

### Subjects

Five healthy adult volunteers (four males, one female) with normal or corrected-to-normal vision participated in this study: S1 (age 32), S2 (age 28), S3 (age 28), S4 (age 28), S5 (age 28). Data were collected at the University of California, Berkeley. The main experiment contained three sessions. Functional localizers were collected in two sessions. The protocols for these experiments were approved by the Committee for the Protection of Human Subjects at the University of California, Berkeley. Written informed consent was obtained from all subjects before scanning.

### Stimuli

Continuous natural movies were used as the stimulus in the main experiment. Three 8 min movies were compiled from short clips. Movie clips were selected from a wide variety of sources as detailed in Nishimoto et al. (2011). High-definition movie frames were cropped into a square frame and downsampled to 512 × 512 pixels, covering a 24°× 24° field of view. Subjects were directed to fixate on a color square of 0.16°× 0.16°at the center. The color of the fixation dot was changing at 1 Hz to ensure visibility. The stimulus was presented at a rate of 15 Hz using an MRI-compatible projector (Avotec) and a custom-built mirror arrangement.

### Experiment design

The main experiment was performed in a single session consisting of nine 8 min runs. Four mutually exclusive classes of stimuli (only humans, only vehicles, both humans and vehicles, and neither humans nor vehicles) were randomly interleaved and evenly distributed within and across the runs. Subjects were instructed to covertly search for the target categories in the movies. A cue word was displayed before each run to indicate the attention task: “humans”, “vehicles”, or “humans and vehicles”. In the attend to humans task, subjects searched for human categories (e.g. woman, man, boy). For the attend to vehicles task, subjects searched for vehicle categories (e.g. car, truck, bus). Subjects searched for targets from either of the human or vehicle categories in the divided attention task. To minimize subject expectation bias, the order of search tasks was interleaved across runs. To maintain vigilance, subjects were asked to press a button when they detected a target on the screen. BOLD responses were recorded from the whole brain. To minimize the effect of transient confounds, data from the first 20 seconds and the last 30 seconds of each run were discarded. These procedures resulted in 690 data samples for each attention task.

### Definition of functional areas

Functional regions of interest (ROIs) were identified in individual subjects using functional localizers (Huth et al., 2012). Localizer experiments for category-selective areas (fusiform face area, FFA; extrastriate body area, EBA; parahippocampal place area, PPA; retrosplenial cortex, RSC; lateral occipital complex, LOC) were performed in six 4.5 min runs of 16 blocks (Huth et al., 2012). Subjects passively viewed 20 random static images from one of the objects, scenes, body parts, faces, or spatially scrambled objects groups in each block. Each image was shown for 300 ms following a 500 ms blank period. Scene-selective ROIs (PPA, RSC) were identified as voxels with positive scene versus objects contrast (*t*-test, *p <* 10^−4^, uncorrected). FFA, EBA, and LOC were defined using face-versus-object, body-part-versus-object and object-versus-scrambled-object contrasts respectively (*t*-test, *p <* 10^−4^, uncorrected). Localizer experiment for attentional-control areas (intraparietal sulcus, IPS; frontal eye field, FEF; frontal operculum, FO) contained one 10 min run. In each 20 sec block, either a self-generated saccade task or a resting task was prescribed (Corbetta et al., 1998). The run contained 30 blocks. Attentional-control areas were localized using saccade-versus-rest contrast (*t*-test, *p <* 10^−4^, uncorrected). ROIs were refined to voxels with contrast level more than half of the maximum near a 2 mm neighborhood of the cortical surface.

### MRI protocols

Data were collected using a 3T Siemens Tim Trio MRI scanner (Siemens Medical Solutions) using a 32-channel receiver coil. Functional data were collected using a T2*-weighted gradient-echo echo-planar-imaging pulse sequence with the following parameters: TR = 2 sec, TE = 33 msec, water-excitation pulse with flip angle = 70°, voxel size = 2.24 mm×2.24 mm×4.13 mm, field of view = 224 mm×224 mm, 32 axial slices. To construct cortical surfaces, anatomical data were collected using a three-dimensional T1-weighted magnetization-prepared rapid-acquisition gradient-echo sequence with the following parameters: TR = 2.3 sec, TE = 3.45 msec, flip angle = 10°, voxel size = 1 mm×1 mm×1 mm, field of view = 256 mm×212 mm×256 mm.

### Data preprocessing

Functional images collected in the main experiment were motion corrected. Using the SPM12 software package (Friston et al., 1995), the functional images were aligned to the first image from the first session of the main experiment. Three subjects that participated in this study were common with a previous study (Çukur et al., 2013). Functional images of these subjects were aligned to the reference images from that previous study. Non-brain tissues were removed using the brain extraction tool (BET) from the FSL software package (Smith, 2002). Within each run, low-frequency drifts were removed from BOLD responses in each voxel using a second order Savitzky-Golay filter over a 240 sec temporal window. The resulting voxelwise time series were z-scored to attain zero mean and unity variance.

### Voxelwise category model

For each voxel, response profiles were determined by fitting category models that represented hundreds of objects and actions in natural movies. As a first step, each 1sec clip of the movie stimulus was manually labeled for presence of 831 distinct object and action categories (Huth et al., 2012). Presence of superordinate categories was inferred from the terms in the WordNet lexicon, a lexical database that groups words based on their semantic relationships (Miller, 1995). This procedure yielded time courses for 831 model features (i.e. categories). Each time course was then downsampled to 0.5 Hz to match the acquisition rate of fMRI. Separate finite impulse response (FIR) filters were used for each model feature to capture the hemodynamic response. Filter delays were set to 4, 6, and 8 secs. This is equivalent to concatenating feature vectors that are delayed by two, three, and four samples.

To prevent head-motion and physiological noise confounds, estimates of these nuisance factors were regressed out of the BOLD responses. Six affine motion time courses estimated during the motion-correction stage were taken as the head-motion regressors. Two regressors to capture respiration and nine regressors to capture cardiac activity were estimated using the data collected via a pulse oximeter and a pneumatic belt during the main experimental runs (Verstynen and Deshpande, 2011).

To reduce spurious correlations between model features and global motion-energy of the movie stimulus, a nuisance regressor was included that reflected the total motion-energy. The motion-energy time course was formed by taking the mean motion-energy in each one second movie clip. Movie frames were transformed into the International Commission on Illumination LAB color space, and the luminance channel was extracted. The luminance was then passed through the motion-energy filter bank. The motion-energy filter bank contained 2139 Gabor filters. Filters were computed at eight directions (0 to 315°, in 45°steps), three temporal frequencies (0, 2, and 4 Hz) and six spatial frequencies (0, 1.5, 3, 6, 12, and 24 cycles/image). Filters were placed on a square grid spanning the 24°× 24°field of view. Finally, the motion-energy time course was assessed by squaring and summing outputs of quadrature filter pairs, and the results were passed through a logarithm compressive nonlinearity and temporally downsampled to match the fMRI acquisition rate (Nishimoto et al., 2011).

To account for potential correlations between target detection and BOLD responses, a target-presence regressor was included in the model. The target-presence regressor contained category regressor for “person” during attend to humans task and the category regressor for “conveyance” during attend to vehicles task. The target-presence regressor during divided attention task contained the binary union of the “person” and “conveyance” category regressors. The described regressors were aggregated and used as the stimulus matrix.

### Model fitting and testing

Voxelwise models were fit using regularized linear regression with an ℓ_2_ penalty to avoid overfitting. To prevent bias, model fitting for the three attention tasks was performed concurrently. To do this, the stimulus and BOLD response matrices were aggregated across tasks (Fig.2). Note that this procedure ensures that the same regularization parameter will be used in each voxel across the three tasks. Furthermore, using the aggregated stimulus matrix enables employing the target regressor.

**Figuer 2.**
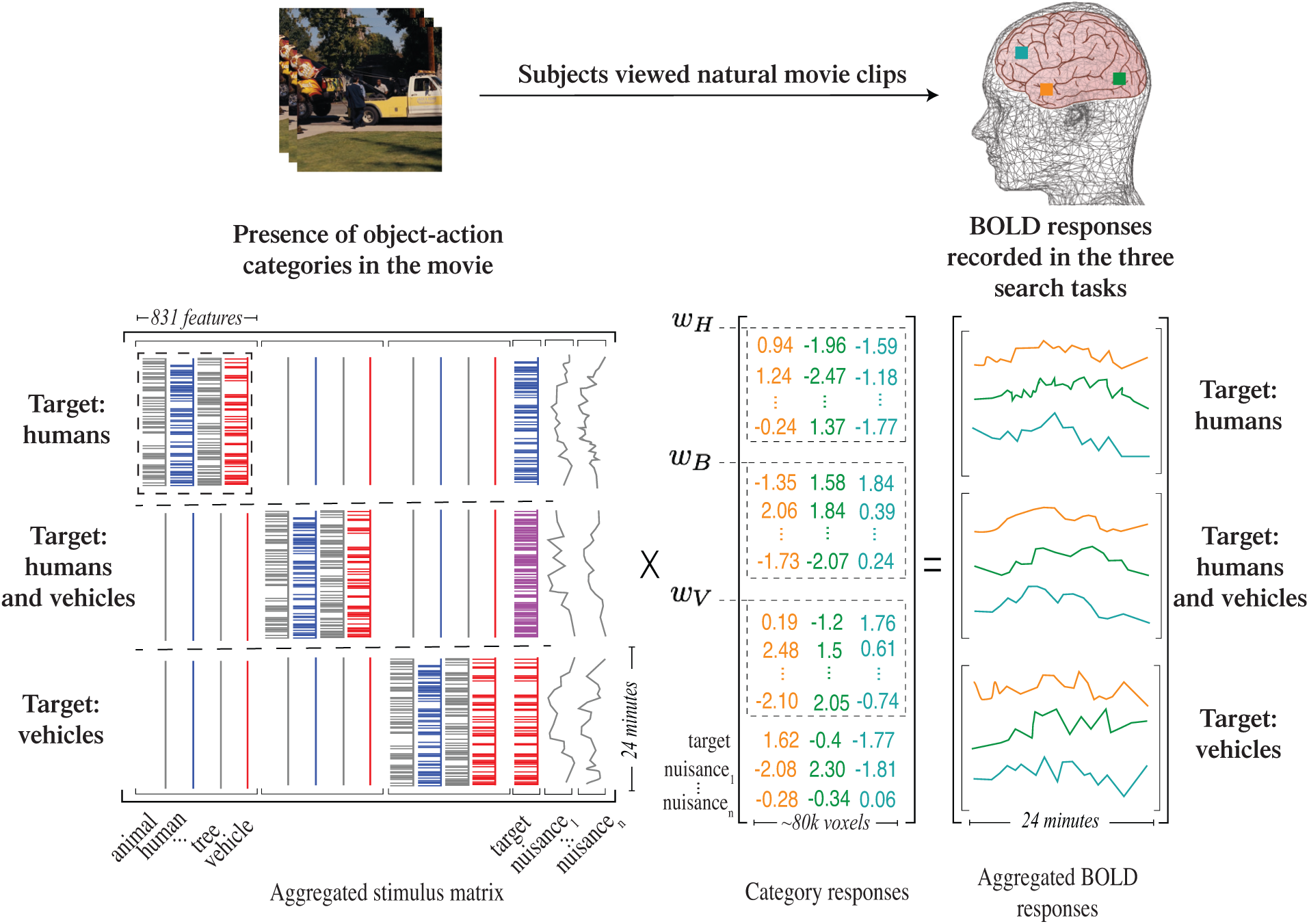
Experimental and modeling procedures. Subjects viewed 72 minutes of natural movies. BOLD responses were recorded using functional MRI (fMRI) while subjects performed covert search for “humans”, “vehicles” (i.e. single-target attention tasks), or “humans and vehicles” (i.e. divided attention task) in the movies. Movies were labeled for presence of object and action categories. Presence of superordinate categories was inferred using the WordNet lexicon that resulted in a total of 831 object and action categories. A target-presence regressor was used to account for BOLD response modulations resulting from detection of targets in the scenes. Target-presence regressor comprised of the human regressor (red series), vehicle regressor (blue series), and the binary union of the two (cyan series) to indicate the presence of “humans”, “vehicles”, and “humans and vehicles”, respectively. Nuisance regressors were used to account for physiological noise. Category models were fit independently for each voxel using regularized linear regression. Category responses represent the contribution of each of 831 object-action categories to BOLD responses. Models for the three attention tasks were fit simultaneously using the aggregated stimulus and BOLD response matrices (*w*_*H*_, *w*_*V*_, and *w*_*B*_ for the attend to “humans”, attend to “vehicles”, and attend to “humans and vehicles” tasks, respectively).

A nested cross-validation (CV) procedure was used to estimate response profiles for each voxel. Data were segmented into 58 25sec blocks. In each of the 20 outer folds, 6 blocks were randomly held-out as validation data and the remaining blocks were used for parameter optimization and fitting models on the inner folds. In each of the 20 inner folds, blocks were randomly shuffled and split to 40 blocks as training data and 12 blocks as test data. Models were fit on the training data for regularization parameters in the range [2^−3^, 2^20^]. Using the weights found for each regularization parameter, responses were predicted for the test data. Prediction scores were separately computed for each voxel, taken as the Pearson’s correlation between actual and predicted responses. Prediction scores were then averaged across the inner CV folds. Regularization parameters maximizing the average prediction score were selected in each voxel. Nuisance regressors were discarded from further analyses. Afterwards, optimized parameters were used to fit models on the union of training and test data in each outer fold, yielding category response profiles. To assess model performance, responses were predicted for the validation data using the fit models and prediction scores of each voxel were averaged across the attention tasks. Finally, response profiles and prediction scores for each voxel were averaged across the outer folds. Model fitting was performed using custom-written software in Matlab (MathWorks MA).

### Significance of the fit models

Significance of the estimated category response profiles was assessed using a bootstrapping procedure on the validation data held-out in the model fitting stage. A 20-fold CV procedure was implemented to assess significance of the fit models. In each fold, the corresponding fit models and validation data from the model fitting stage were used. Responses were predicted for the validation data using the fit models. Prediction score was calculated in each fold and averaged across CV folds to get the mean score. In each fold, predicted responses were resampled 500 times with replacement. We assumed the null hypothesis that attention does not alter response profiles of voxels. Thus, under the null hypothesis, response profiles would be the same across the three attention tasks (Çukur et al., 2013). To get the prediction score distribution under the null hypothesis, resampled predicted responses were shuffled across the attention tasks, and prediction scores were computed. The *p*-value was taken as the fraction of samples for which the average prediction score across CV folds was lower than that under the null hypothesis.

### Semantic representation of objects and actions

To obtain a quantitative description of the individual subjects’ semantic spaces, principal component analysis (PCA) was performed on estimated response profiles in individual subjects (Huth et al., 2012). To prevent bias, PCA was performed on response profiles for the two single-target attention tasks pooled together. The collection of principal components (PCs) that described at least 90% of the variance in response profiles was selected. This resulted in *L* ∈ [36, 47] PCs for the five subjects. The semantic representation of individual object or action categories can then be assessed in individual subjects by projecting the response profiles onto the PCs.

### Linearity of semantic tuning during divided attention

To test whether semantic tuning during divided attention could be predicted using a weighted linear combination of semantic tuning during the two single-target tasks, we compared voxelwise semantic tuning across search tasks. A mask vector, 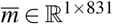, was used to select the categories of interest among 831 categories. Elements of 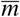 were one for categories of interest and zero elsewhere. Masked response profiles, 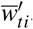, were obtained by element-wise multiplication

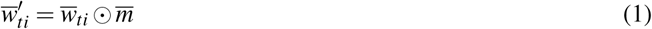

where 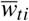 is the response profile for voxel *i* and task *t* ∈ {*H,V, B*} denoting attend to “humans”, attend to “vehicles”, and attend to “both humans and vehicles” (Fig.2), and ⊙ represents element-wise multiplication. Masked response profiles were then projected onto the PCs to assess semantic tuning profiles, 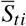

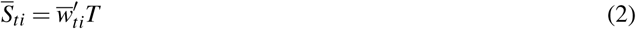

where *T* ∈ ℝ^831×*L*^ is the matrix of *L* PCs. Semantic tuning profile during divided attention was predicted as a weighted linear combination of semantic tuning during the two single-target tasks using ordinary least-squares. A voxelwise linearity index (LI) was then quantified as the Pearson’s correlation coefficient between measured and predicted semantic tuning during divided attention (*Ŝ*_*Bi*_; Fig.4a)

**Figuer 3.**
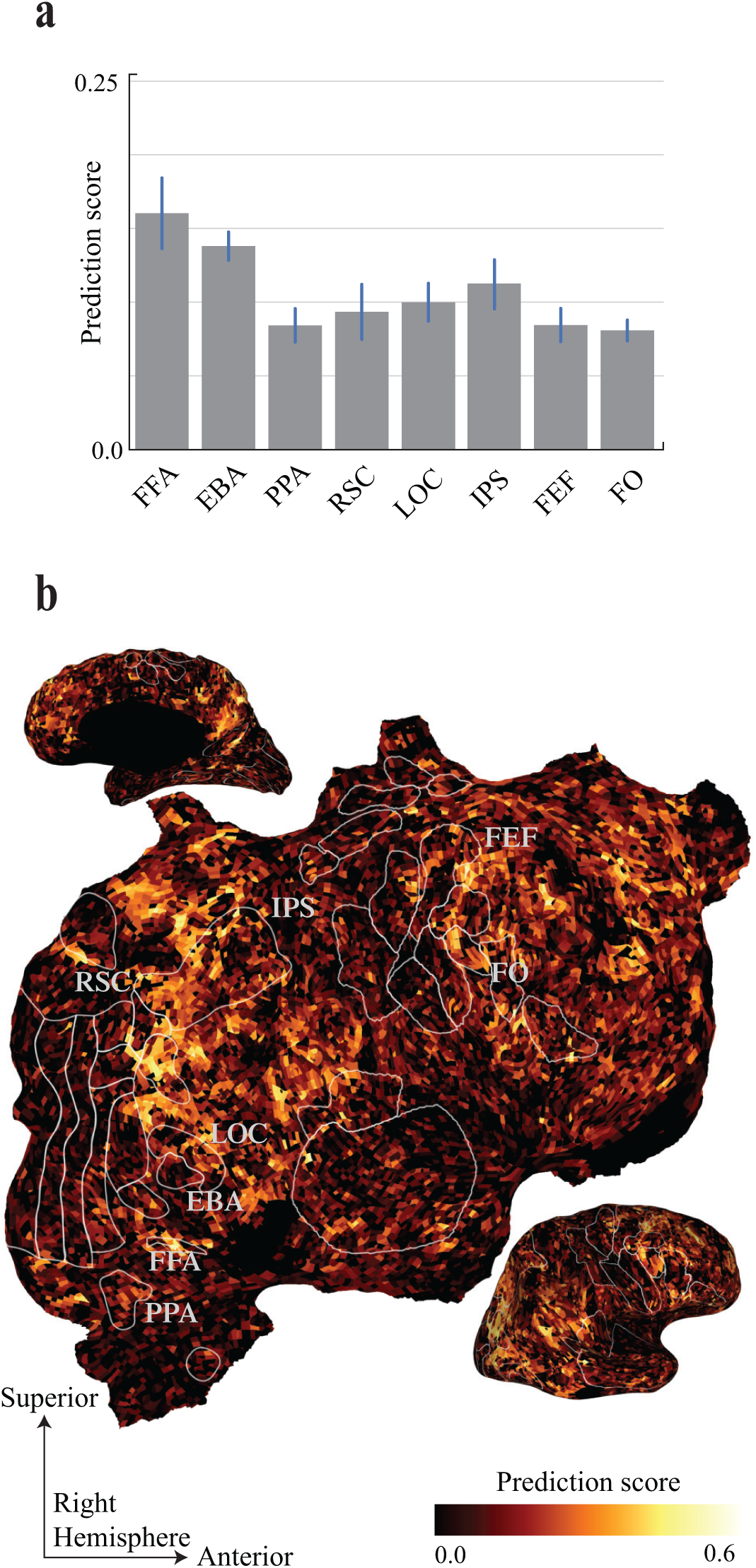
Prediction performance of the category model. To assess the performance of the category model, BOLD responses were predicted using the estimated category responses in each voxel. Pearson’s correlation coefficient between the predicted and measured BOLD responses was taken as the prediction score. **(a)** Prediction score in functional cortical areas (mean*±*s.e.m. across five subjects). FFA, fusiform face area; EBA, extrastriate body area; PPA, parahippocampal place area; RSC, retrosplenial cortex; LOC, lateral occipital complex; IPS, intraparietal sulcus; FEF, frontal eye fields; FO, frontal operculum. **(b)** Cortical flat map of the prediction score for a representative subject. Prediction scores are shown in the right hemisphere. Voxels with high prediction scores appear in yellow color and voxels that have low prediction scores appear in dark gray color. Most voxels in the occipitotemporal cortex, parietal cortex, and prefrontal cortex are well modeled.

**Figuer 4.**
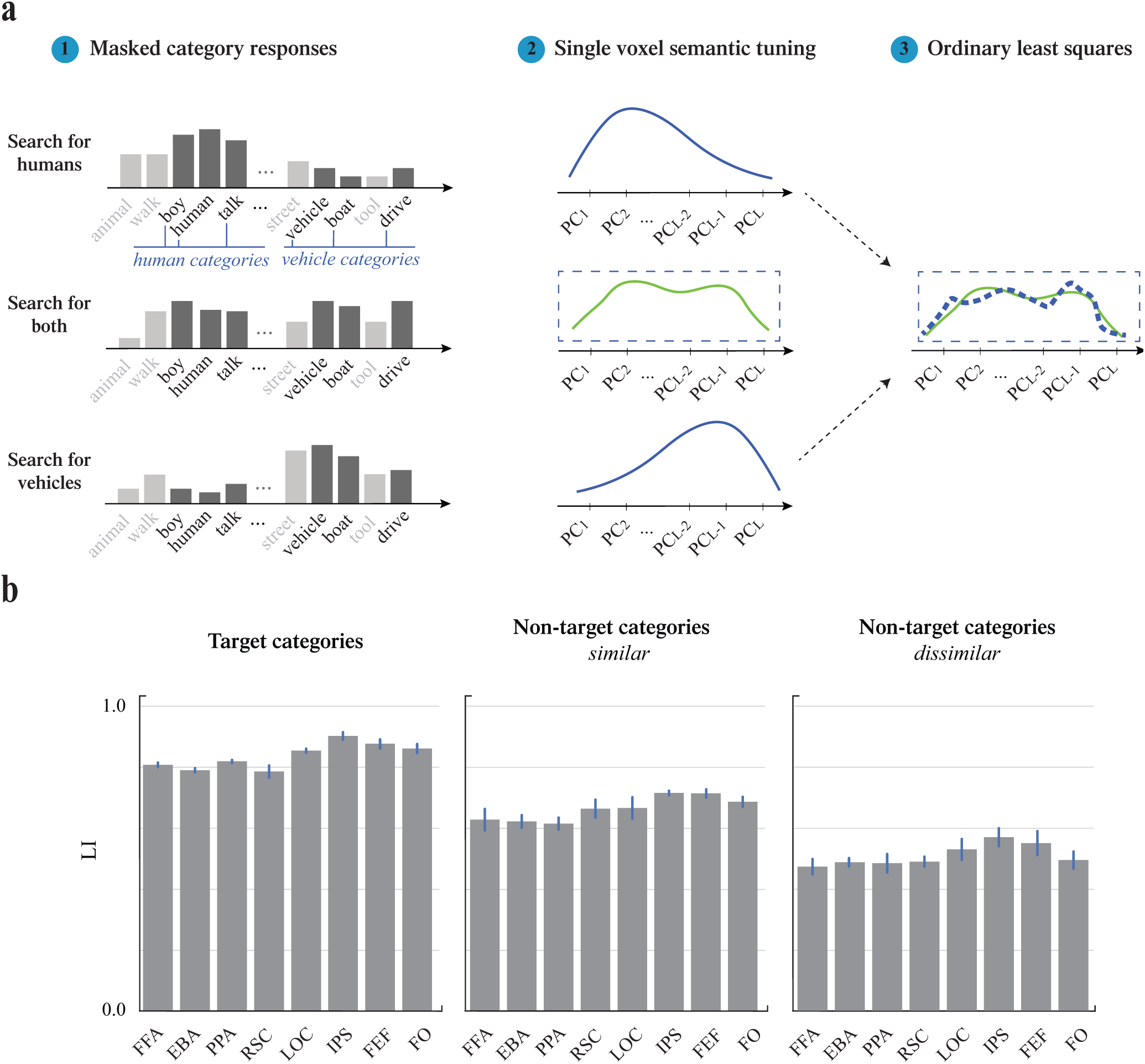
Linearity of semantic tuning during divided attention. **(a)** Masked category responses for the three attention tasks were projected onto subjects’ semantic spaces to estimate the voxelwise semantic tuning profiles. The semantic spaces were estimated by performing principal component analysis (PCA) on response profiles pooled during the two single target attention tasks in individual subjects. The collection of principal components (PCs) explaining at least 90% of the variance in the data was selected. Semantic tuning profile during divided attention was predicted via a weighted linear combination of tuning profiles during the two single target tasks (dashed lines indicate predicted tuning; solid lines indicate measured tuning). Pearson’s correlation coefficient between the predicted and measured semantic tuning during divided attention was taken as the linearity index (LI). **(b)** LI in functional cortical areas (mean*±*s.e.m. across five subjects) for target categories **(left)**, nontarget categories that are semantically similar to targets **(middle)**, and nontarget categories that are dissimilar to targets **(right)**. A substantial portion of semantic tuning during divided attention is described as a weighted linear combination even in the absence of target categories. For all cases, LI is significantly higher in attentional-control areas (IPS, FEF and FO; *p* = 0.004 for target categories, *p* = 0.005 for similar nontarget categories, *p* = 0.036 for dissimilar nontarget categories) and in LOC (*p* = 0.023 for target categories, *p* = 0.048 for similar nontarget categories, *p* = 0.001 for dissimilar nontarget categories) than in category-selective areas (FFA, EBA, PPA, and RSC).

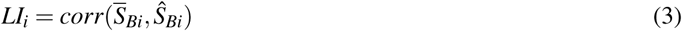

An LI of 1 indicates that semantic tuning during divided attention can be completely described as a weighted linear combination of the tuning profiles during the two single-target tasks. Whereas, an LI of 0 means that semantic tuning during divided attention can not be described linearly in terms of tuning profiles during the two single-target tasks. To study the linearity of the semantic representation during divided attention in an ROI, LIs were averaged across voxels with significant prediction scores within the ROI.

### Bias in semantic representation during divided attention

We questioned whether semantic representation during the divided attention task was biased toward any of the single-target attention tasks. To address this issue, we studied the distribution of semantic representation within an ROI for each individual task. Semantic tuning profiles of significantly predicted voxels within each ROI were pooled to obtain the distribution of tuning profiles

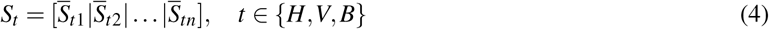

where *S*_*H*_, *S*_*V*_, *S*_*B*_ represent distribution of tuning profiles for attend to “humans”, attend to “vehicles”, and attend to “both humans and vehicles” tasks, and *n* is the number of significantly predicted voxels within the ROI. Note that *S*_*t*_ can also be expressed as

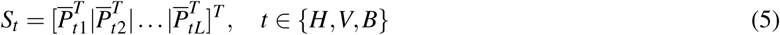

where 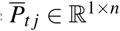 is a row vector that represents the projections of the response profiles for task *t* ∈ {*H,V, B*} on the j^th^ PC across ROI voxels. To emphasize semantic axes that explain higher variance, projections 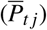 were weighted by the explained variance of the corresponding PCs. This yielded the semantic tuning distribution, 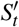 The tuning distribution during divided attention was then regressed onto the distributions during the two single-target tasks

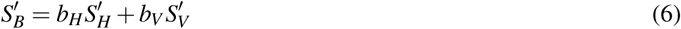

The bias index (BI) was quantified (Fig.5a) as

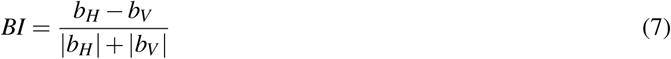

**Figuer 5.**
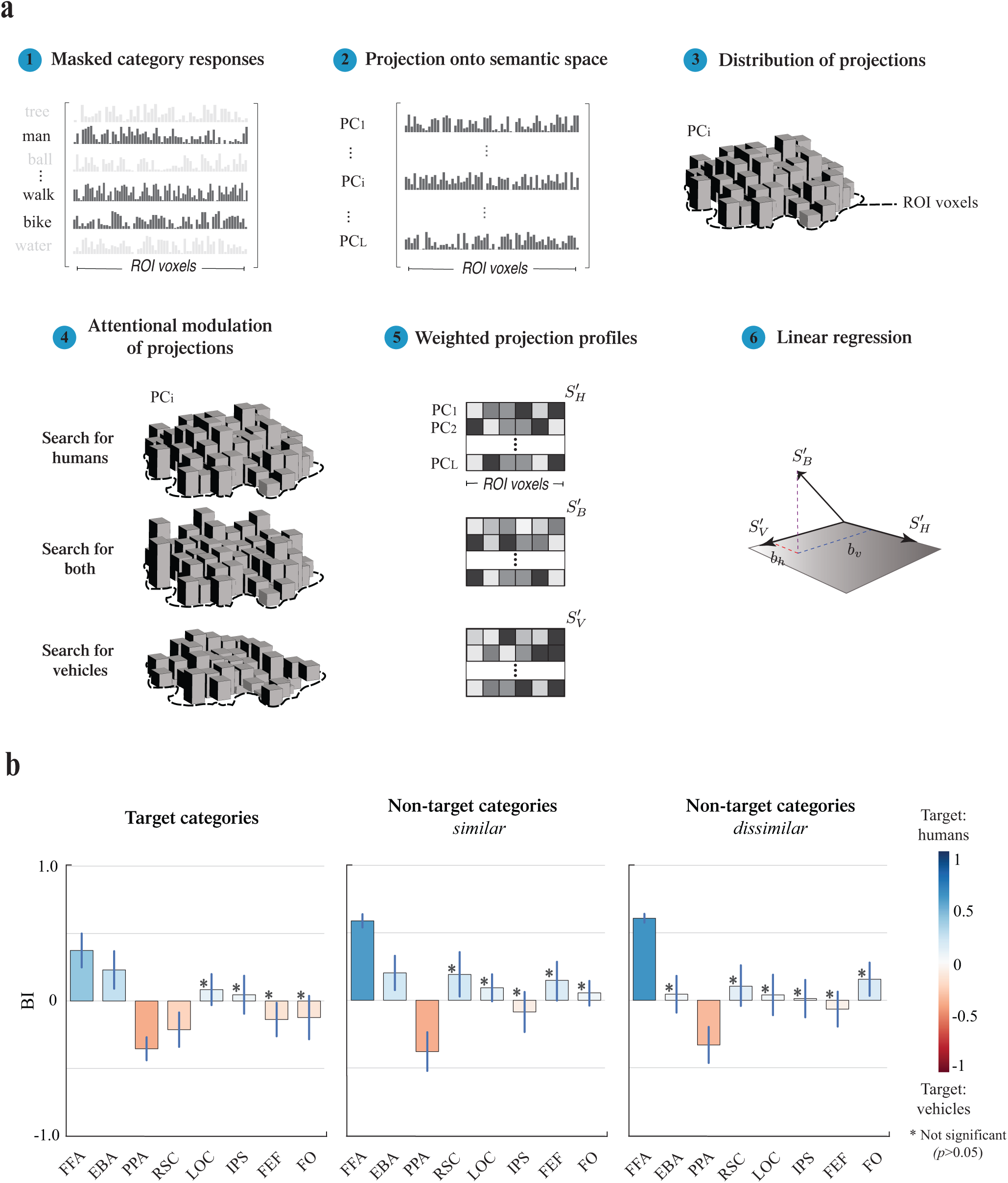
Bias in semantic representation during divided attention. **(a)** To assess the bias in semantic representation during divided attention, masked category responses for the three attention tasks were projected onto semantic spaces in individual subjects. Projections were weighted by the explained variance of PCs to yield the semantic tuning distributions. Semantic tuning distribution during divided attention was then regressed onto tuning distributions during the two single target tasks. Regression weights were used to calculate a bias index (BI). **(b)** BI in functional cortical areas (mean*±*s.e.m. across five subjects) for target categories **(left)**, nontarget categories that are similar to targets **(middle)**, and nontarget categories that are dissimilar to targets **(right)**. Blue versus red bars indicate bias toward attend to humans versus attend to vehicles tasks. Semantic representation of target categories in category-selective areas (FFA, EBA, PPA, and RSC) is biased toward the preferred object-category. Bias is non-significant in attentional-control areas (IPS, FEF and FO) and general object-selective area LOC (bootstrap test, *p >* 0.05). Representation of nontarget categories is also biased toward the preferred target in FFA and PPA.

**Figuer 6.**
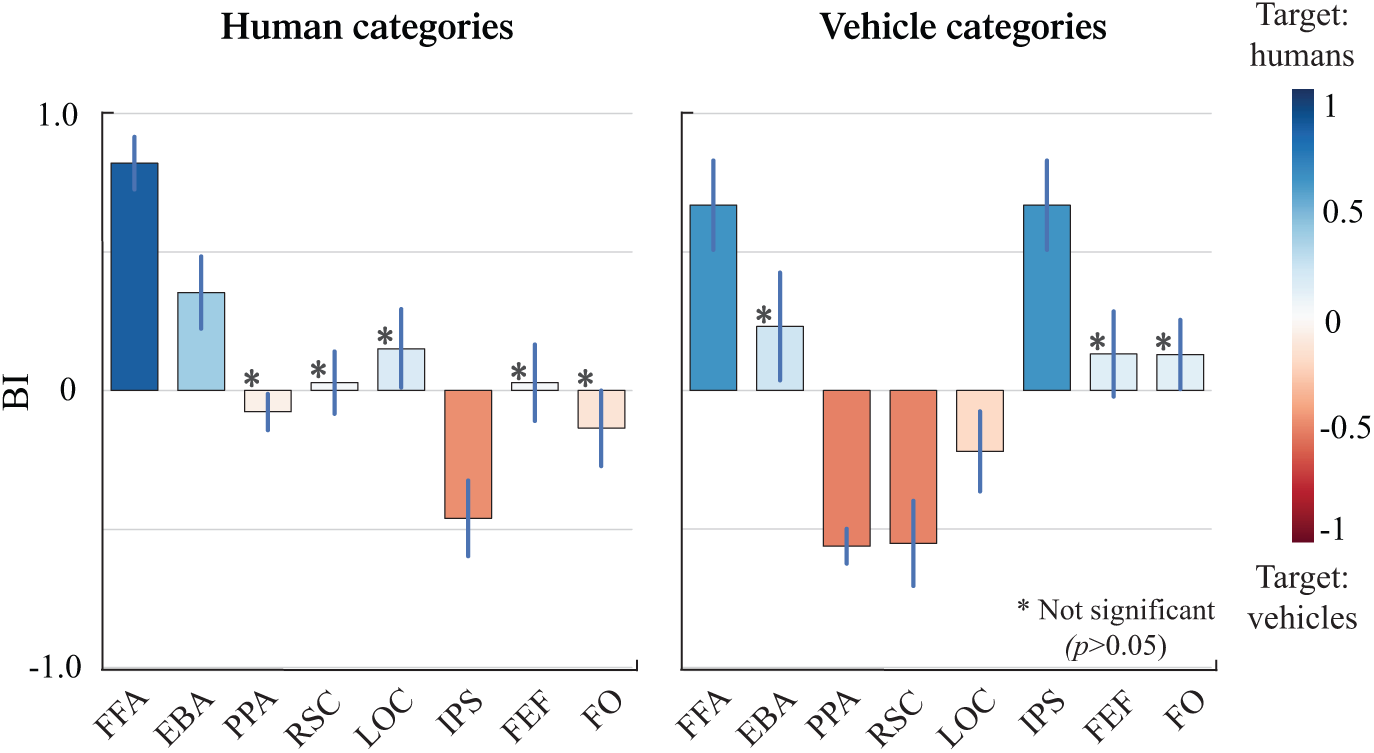
Bias in representation of human and vehicle categories. BI for human categories **(left)**, and vehicle categories **(right)** in functional cortical areas (mean*±*s.e.m. across five subjects). Representations of both human and vehicle categories in FFA are biased toward the attend to humans task. Meanwhile, representation of vehicles but not humans is biased toward the attend to vehicles task in PPA. Representations of both of the target categories are not significantly biased in attentional-control areas in prefrontal cortex. Representation in IPS is biased toward the distractor category.

Bias toward the attend to humans task would yield *BI* ∈ (0, 1]. A BI of 0 means that the tuning distribution during divided attention is not biased toward any of the single-target attention tasks. Whereas, a BI of 1 means that tuning distribution during divided attention is completely biased toward attend to humans task. Similarly, bias toward the attend to vehicles task would yield *BI* ∈ [−1, 0) where a BI of −1 means that tuning during distribution divided attention is completely biased toward attend to vehicles task. Note that since the response profiles for the three tasks were projected onto the same PCs the calculated BI is immune to changes in the direction of PCs.

### Visualization on cortical surfaces

Cortical flatmaps were generated by projecting voxelwise results on individual subject flattened cortical surfaces. Cortical surfaces were constructed for each subject using T1-weighted anatomical brain scans. Freesurfer software was used to construct surfaces (Reuter et al., 2012). Surfaces were then flattened using Pycortex (Gao et al., 2015).

## Results

### Representation of categories during visual search

A recent study showed that thousands of object and action categories in natural scenes are embedded in a semantic space the axes of which are mapped continuously across the cortex (Huth et al., 2012). Moreover, in a previous study from our lab we reported that category-based attention warps this space to expand the representation of targets (Çukur et al., 2013). Yet, attentional modulation of semantic representations during attention to multiple targets remains understudied. To investigate this issue, we estimated voxelwise tuning for hundreds of object and action categories across neocortex. Five human subjects viewed 72 minutes of natural movies while they attended to “humans”, “vehicles” (i.e. single-target attention tasks), or “both humans and vehicles” (i.e. divided attention task) in separate runs. Separate models were fit for each voxel and attention task. This enabled us to measure responses to 831 distinct object and action categories in the three attention tasks (Fig.2, see *Materials and Methods*). We find that the category model accurately predicts responses in many voxels across ventral-temporal, parietal and prefrontal cortices (Fig.3).

We hypothesized that if the category responses get modulated by attention task, the fit models during the three attention tasks would predict better than a null model fit by pooling data across tasks. To investigate this issue, we compared the average prediction score in the three attention tasks with that of a null model. To obtain the null model, predicted responses were shuffled across tasks and prediction scores were computed afterwards. We found that attention significantly modulates category responses in 77.03 *±* 5.47% of cortical voxels (mean*±*s.d. across five subjects; bootstrap test, *p <* 0.05). This result implies that attention modulates category responses across visual and non-visual areas.

### Linearity of semantic tuning during divided attention

We hypothesized that if the attentional modulations during search for multiple targets are mediated by BC, semantic tuning profile during divided attention should be a weighted linear combination of tuning profiles during isolated attention to individual targets. To test this prediction, we projected category response profiles onto individual subjects’ semantic spaces. The semantic space was constructed via principal component analysis (PCA) on voxelwise response profiles pooled across the single-target attention tasks. A collection of 36-47 PCs that accounted for over 90% of the variance were selected. To obtain semantic representations of specific categories, the categories of interest were masked in the response profiles. Ordinary least-squares was then used to predict voxelwise semantic tuning profile during divided attention as a weighted linear combination of semantic tuning profiles during the two single-target attention tasks. Linearity index (LI) was taken as Pearson’s correlation coefficient between the predicted and measured semantic tuning during divided attention (Fig.4a, see *Materials and Methods*). We find that LI for target categories (union of human and vehicle categories) is 0.81 *±* 0.02 in category-selective areas (FFA, PPA, EBA, and RSC; mean*±*s.d.), 0.86 *±* 0.02 in general object-selective area LOC (mean*±*s.e.m), and 0.88 *±* 0.02 in attentional-control areas (IPS, FEF, and FO; mean*±*s.d.; Fig.4b). LI is significantly greater than zero in all of the studied functional areas (bootstrap test, *p <* 10^−4^). These results imply that a substantial portion of semantic tuning during divided attention is linearly described as a weighted linear combination of tuning during attention to individual targets. In LOC, LI is significantly higher than that of category-selective areas (bootstrap test, *p* = 0.023). Moreover, LI in attentional-control areas is significantly higher than that of category-selective areas (bootstrap test, *p* = 0.004). These results suggest that semantic tuning for target categories better conform to the weighted linear combination model in general object-selective area and in later stages of visual processing compared to visual areas that have strong category preference.

A recent study from our laboratory showed that during category-based attention voxelwise tuning for nontarget categories that are semantically similar to targets shifts toward target categories (Çukur et al., 2013). We thus asked whether the prediction performance of the weighted linear combination for nontarget categories that are semantically similar to targets is similar to that for targets. To answer this question, we calculated LI separately for nontarget categories that are semantically similar to targets (i.e. animals, social places, devices, and buildings), and for non-target categories that are dissimilar to targets (all categories except the union of animals, devices, buildings, social places, and target categories). LI for similar categories is 0.64 *±* 0.02 in category-selective areas, 0.67 *±* 0.05 in LOC, and 0.71 *±* 0.02 in attentional-control areas. LI for dissimilar categories is 0.50 *±* 0.01 in category-selective areas, 0.54 *±* 0.06 in LOC, and 0.56 *±* 0.03 in attentional-control areas. LI for similar categories is higher than that for dissimilar categories in all functional ROIs (bootstrap test, *p <* 10^−4^). This finding suggests that during divided attention to multiple targets, the competition in representation of nontarget visual objects is carried over to objects that are similar to targets (McMains and Kastner, 2010; Beck and Kastner, 2007). LI is significantly lower in category-selective areas than in LOC (bootstrap test, *p* = 0.048 for similar categories, *p* = 0.001 for dissimilar categories), and in attentional-control areas (bootstrap test, *p* = 0.007 for similar categories, *p* = 0.019 for dissimilar categories). This result implies that similar to target categories, semantic tuning for nontarget categories is more linear in LOC and in later stages of visual processing compared to visual areas that have strong category preference.

### Bias in semantic representation during divided attention

BC observes that inherent selectivity of cortical areas during passive viewing can bias the competition in favor of the preferred target during visual search (Desimone and Duncan, 1995). To study the interactions between the attentional bias in semantic representation and the intrinsic category-selectivity of cortical areas, we expressed the semantic representation during divided attention task as a weighted linear combination of the semantic representations during the two single-target tasks in individual cortical areas. We then investigated whether weights in the weighted linear combination were biased toward any of the single-target attention tasks. Masked response profiles across voxels within the ROI were projected onto individual subjects’ semantic spaces to assess the semantic tuning distribution. We regressed the semantic tuning distribution during divided attention onto distributions during the two single-target tasks. We then quantified a bias index (BI) using the regression weights. According to this index, bias in the semantic representation during divided attention task toward the attend to humans task versus attend to vehicles task was represented as positive versus negative BI in the range [−1, 1] (Fig.5a, see *Materials and Methods*). We find that BI for target categories is 0.32 *±* 0.09 in human-selective areas (FFA and EBA), and −0.29 *±* 0.12 in scene-selective areas (PPA and RSC; mean*±*s.d.; bootstrap test, *p <* 10^−4^; Fig.5b). BI is non-significant in attentional-control areas (IPS, FEF, FO) and the general object-selective area LOC (bootstrap test, *p >* 0.05). These results suggest that the competition in representation of target categories during divided attention is biased in favor of the preferred target in cortical areas that are strongly selective for targets. On the contrary, semantic representation is not biased in the areas without any specific category preference.

In a previous study we showed that attention shifts semantic tuning for both target and nontarget categories (Çukur et al., 2013). Thus, we asked if there is any bias in representation of nontarget categories during divided attention. To answer this question we separately calculated BI for nontarget categories that are similar to targets and nontarget categories that are dissimilar to targets. BI for similar categories is 0.38 *±* 0.25 in human-selective areas, and −0.11 *±* 0.38 in scene-selective areas (mean*±*std.; bootstrap test, *p <* 10^−4^; non-significant in RSC (*p* = 0.288)). BI is non-significant in attentional-control areas and in LOC (bootstrap test, *p >* 0.05). Meanwhile, BI for nontarget dissimilar categories is 0.32 *±* 0.37 in human-selective areas, and −0.11 *±* 0.27 in scene-selective areas (mean*±*std.; bootstrap test, *p <* 0.05; non-significant in EBA (*p* = 0.610) and in RSC (*p* = 0.754)). BI is non-significant in attentional-control areas and in LOC (bootstrap test, *p >* 0.05). These results indicate that representation of nontarget categories that are similar to targets is biased in favor of the preferred target in category-selective areas. Yet, representation of nontarget categories that are dissimilar to targets is only biased in areas that are strongly selective for the targets and not in areas that are selective for categories that are semantically similar to targets.

The target categories used here (i.e. humans and vehicles) show high semantic dissimilarity. This raises the possibility that the biases in semantic representation differ between human categories and vehicle categories. To examine this issue, we compared BI for human and vehicle categories separately. We find that BI for human categories is 0.59 *±* 0.31 in human-selective areas, and −0.02 *±* 0.08 in scene-selective areas (mean*±*std.; bootstrap test, *p <* 10^−4^ in FFA, EBA; non-significant in PPA (*p* = 0.88), and in RSC (*p* = 0.94)). Whereas, BI for vehicle categories is 0.38 *±* 0.29 in human-selective areas, and −0.50 *±* 0.01 in scene-selective areas (mean*±*std.; bootstrap test, *p <* 10^−4^; non-significant in EBA (*p* = 0.62)). BI in human-selective areas is significantly positive for both human categories and vehicle categories. Yet, it is stronger for human categories compared to vehicle categories. Whereas, in scene-selective areas, bias is significant only for vehicle categories. These results imply that scene-selective areas have a more dynamic representation of their nonpreferred object categories compared to human-selective areas during divided attention.

Among the attentional-control areas, BI is significant for both human and vehicle categories only in IPS (bootstrap test, *p <* 10^−4^). In IPS, BI is −0.48 *±* 0.13 for human categories, and 0.60 *±* 0.14 for vehicle categories (mean*±*s.e.m). This result indicates that in IPS, representation of human categories is biased toward vehicles and representation of vehicle categories is biased toward humans. Previous studies suggest that IPS is involved in neural circuits that enhance detection of distractors (Greenberg et al., 2010; Mevorach et al., 2010; Sakai et al., 2002; Bledowski et al., 2004). In line with these studies, this finding suggests that IPS is involved in category-based attention by enhancing the representation of distractors.

## Discussion

In this study, we tested whether the biased competition hypothesis can account for modulation of semantic representations during a divided attention task. We fit a category model to characterize category responses of single voxels during search for “humans”, “vehicles”, and “both humans and vehicles”. We found that the category model explains significant response variance in many voxels across ventral-temporal, parietal, and prefrontal cortices. We estimated the semantic space underlying category models, and then assessed semantic representations by projecting the category responses for the three search tasks onto the semantic space.

### Linearity of the semantic representation during divided attention

We find that a large portion of the variance in semantic tuning during divided attention can be explained using a weighted linear combination of tuning during isolated attention to individual targets. We find that semantic tuning for target categories is more accurately predicted via the weighted linear combination relationship compared to semantic tuning for nontarget categories. In a recent study, we reported that attention shifts semantic tuning for target categories to a higher degree compared to that for nontarget categories (Çukur et al., 2013). Thus, our results can be attributed to the higher degree of attentional tuning shift for target categories compared to that for nontarget categories.

Several previous studies have investigated differences in the level of competition between strongly category-selective areas and areas without specific category preference. Reddy and Kanwisher (2007) and MacEvoy and Epstein (2009) showed that response patterns to a pair of objects can be better predicted by a linear combination of responses to constituent objects in LOC compared to in FFA or PPA. In line with these studies, here we find that semantic representation is more linear in LOC compared to that in category-selective areas. These results raise the possibility that semantic representation may also be more linear in attentional-control areas which are not selective for any specific categories compared to strongly category-selective areas (Çukur et al., 2013). Here we find that semantic representations of target and nontarget categories during divided attention are better explained using the weighted linear combination of representations during search for individual targets in attentional-control areas compared to category-selective areas. Our results suggest that higher order areas that are not tuned for specific categories have a more flexible representation of natural scenes during divided attention.

### Bias in semantic representation during divided attention

#### Category-selective areas

The bias in the competition among representation of multiple objects have been investigated in several previous studies. Reddy et al. (2009) reported that BOLD responses to multiple static images of faces, houses, shoes, or cars in category-selective areas, FFA and PPA, are biased toward the preferred target. In line with this finding, here we find that semantic representation of target categories during divided attention in category selective areas FFA, EBA, RSC, and PPA is biased toward the preferred category. We have previously shown that category-based attention shifts semantic tuning not only for the target, but also for nontarget categories that are semantically similar to the target (Çukur et al., 2013). Consistent with this view, here we find that semantic representations of nontarget categories that are similar to targets in FFA and PPA are also biased toward the preferred target. Furthermore, representation of nontarget categories that are dissimilar to targets is biased in FFA and PPA, albeit to a lower degree. This finding suggests that in category-selective areas, semantic similarity to targets enhances the level of bias in semantic representation.

In human-selective areas, we find that the semantic representations of target categories (both humans and vehicles) during divided attention is biased toward the representation during the attend to humans task. Meanwhile, in vehicle-selective areas, the representation of vehicles but not humans is biased toward the representation during the attend to vehicles task. This suggests a differential role for human- and vehicle-selective areas in representing nonpreferred targets during divided attention. A potential explanation of this result is that vehicle-selective areas have a more dynamic representation of nonpreferred targets compared to human-selective areas (Grill-Spector et al., 2004).

#### Attentional-control areas

We did not observe bias in semantic representation of targets toward any of the target categories in the prefrontal areas that are considered to be part of the attentional-control network. This is expected considering the lack of tuning to specific categories in these areas (Huth et al., 2012). In IPS, we find that semantic representation of humans during divided attention is biased toward the representation during the attend to vehicles task and that the representation of vehicles during divided attention is biased toward the representation during the attend to humans task. Several previous studies suggest that areas in parietal cortex including IPS enhance visual search by maintaining the representation of distractors (Mevorach et al., 2010; Bledowski et al., 2004), in addition to spatial guidance of attention toward targets (Ptak, 2012; Preston et al., 2013). Consistent with this hypothesis, our results can be interpreted to imply that IPS facilitates natural visual search by maintaining representations of distractor categories.

### Limitations and future work

Natural stimuli contain correlations among various levels of features (Hamilton and Huth, 2018). For instance, there might be correlations among low-level visual features of natural scenes and object categories within these scenes (Lescroart et al., 2015). Such correlations can then bias the category responses that we estimated here, which lead to a biased assessment of semantic representations. To minimize correlations between category features and global motion-energy of the movie clips, we used a motion-energy regressor in our modeling procedure. However, we do not rule out the possibility that there might be residual correlations between category features and intermediate features of the movie stimuli, such as object shape characteristics (Op de Beeck et al., 2008) and scene layout (Mullally and Maguire, 2011). Note that it is perhaps impossible to create natural stimuli completely free from these correlations. However, future studies may mitigate this problem by compiling more controlled natural stimuli that minimize stimulus correlations while maintaining high variance in individual categories.

Here we examined weighted linear combination as a competition model for the attentional modulation of semantic representation during divided attention. However, nonlinear models of competition have also been proposed for single neuron responses in primates. Heuer and Britten (2002) found that in macaque area MT, responses to a pair of Gabor patches with high versus low contrast was similar to the responses to the Gabor patch with high contrast. Gawne and Martin (2002) showed that in macaque area V4, responses to a pair of checkerboard patterns is similar to the maximum of the responses to individual patterns in the pair. Further work is needed to investigate possible nonlinear models of competition in semantic representations.

In conclusion, our results imply the presence of biased competition not only at the level of individual objects but also at the level of high-level semantic representations. This competition is evident not only among targets but also among nontarget categories in natural scenes. Yet, the linearity and bias in representation of nontargets depend on their semantic similarity to targets. Overall, these results help explain the human ability to perform concurrent search for multiple targets in complex visual environments.

## Acknowledgements

This work was supported in part by a Marie Curie Actions Career Integration Grant (PCIG13-GA-2013-618101), by a European Molecular Biology Organization Installation Grant (IG 3028), by a TUBA GEBIP 2015 fellowship, and by a BAGEP 2017 fellowship.

## References

Baeck A, Wagemans J, Op de Beeck HP (2013) The distributed representation of random and meaningful object pairs in human occipitotemporal cortex: The weighted average as a general rule. NeuroImage 70:37–47.

Beck DM, Kastner S (2007) Stimulus similarity modulates competitive interactions in human visual cortex. Journal of Vision 7:19.1–12.

Bichot NP, Rossi AF, Desimone R (2005) Parallel and Serial Neural Mechanisms for Visual Search in Macaque Area V4. Science 308:529–534.

Bledowski C, Prvulovic D, Goebel R, Zanella FE, Linden DEJ (2004) Attentional systems in target and distractor processing: a combined ERP and fMRI study. NeuroImage 22:530–540.

Boynton GM (2005) Attention and visual perception. Current Opinion in Neurobiology 15:465–469.

Corbetta M, Akbudak E, Conturo TE, Snyder AZ, Ollinger JM, Drury HA, Linenweber MR, Petersen SE, Raichle ME, Van Essen DC, Shulman GL (1998) A common network of functional areas for attention and eye movements. Neuron 21:761–773.

Çukur T, Nishimoto S, Huth AG, Gallant JL (2013) Attention during natural vision warps semantic representation across the human brain. Nature Neuroscience 16:763–770.

Desimone R (1998) Visual attention mediated by biased competition in extrastriate visual cortex. Philosophical Transactions of the Royal Society B: Biological Sciences 353:1245–1255.

Desimone R, Duncan J (1995) Neural mechanisms of selective visual attention. Annual Review of Neuroscience 18:193–222.

Duncan J (1984) Selective attention and the organization of visual information. Journal of Experimental Psychology: General 113:501–517.

Eckstein MP, Thomas JP, Palmer J, Shimozaki SS (2000) A signal detection model predicts the effects of set size on visual search accuracy for feature, conjunction, triple conjunction, and disjunction displays. Perception & Psychophysics 62:425–451.

Friston KJ, Frith CD, Turner R, Frackowiak RSJ (1995) Characterizing evoked hemodynamics with fMRI. NeuroImage 2:157–165.

Gao JS, Huth AG, Lescroart MD, Gallant JL (2015) Pycortex: an interactive surface visualizer for fMRI. Frontiers in Neuroinformatics 9:162.

Gawne TJ, Martin JM (2002) Responses of Primate Visual Cortical V4 Neurons to Simultaneously Presented Stimuli. Journal of Neurophysiology 88:1128–1135.

Gentile F, Jansma BM (2010) Neural competition through visual similarity in face selection. Brain Re-search 1351:172–184.

Greenberg AS, Esterman M, Wilson D, Serences JT, Yantis S (2010) Control of spatial and feature-based attention in frontoparietal cortex. The Journal of Neuroscience 30:14330–14339.

Grill-Spector K, Knouf N, Kanwisher NG (2004) The fusiform face area subserves face perception, not generic within-category identification. Nature Neuroscience 7:555–562.

Hamilton LS, Huth AG (2018) The revolution will not be controlled: natural stimuli in speech neuroscience. Lan-guage, Cognition and Neuroscience 27:1–10.

Heuer HW, Britten KH (2002) Contrast dependence of response normalization in area MT of the rhesus macaque. Journal of Neurophysiology 88:3398–3408.

Huth AG, Nishimoto S, Vu AT, Gallant JL (2012) A continuous semantic space describes the representation of thousands of object and action categories across the human brain. Neuron 76:1210–1224.

Jeong SK, Xu Y (2017) Task-context-dependent Linear Representation of Multiple Visual Objects in Human Parietal Cortex. Journal of Cognitive Neuroscience 29:1778–1789.

Kastner S, De Weerd P, Desimone R, Ungerleider LG (1998) Mechanisms of directed attention in the human extrastriate cortex as revealed by functional MRI. Science 282:108–111.

Keitel C, Andersen SK, Quigley C, Müller MM (2013) Independent Effects of Attentional Gain Control and Competitive Interactions on Visual Stimulus Processing. Cerebral Cortex 23:940–946.

Lescroart MD, Stansbury DE, Gallant JL (2015) Fourier power, subjective distance, and object categories all provide plausible models of BOLD responses in scene-selective visual areas. Frontiers in Computational Neuroscience 9:1–20.

Luck SJ, Chelazzi L, Hillyard SA, Desimone R (1997) Neural Mechanisms of Spatial Selective Attention in Areas V1, V2, and V4 of Macaque Visual Cortex. Journal of Neurophysiology 77:24–42.

MacEvoy SP, Epstein RA (2009) Decoding the representation of multiple simultaneous objects in human occipitotemporal cortex. Current Biology 19:943–947.

McMains S, Kastner S (2011) Interactions of top-down and bottom-up mechanisms in human visual cortex. The Journal of Neuroscience 31:587–597.

McMains SA, Kastner S (2010) Defining the Units of Competition: Influences of Perceptual Organization on Competitive Interactions in Human Visual Cortex. Journal of Cognitive Neuroscience 22:2417–2426.

Mevorach C, Hodsoll J, Allen H, Shalev L, Humphreys G (2010) Ignoring the elephant in the room: a neural circuit to downregulate salience. The Journal of Neuroscience 30:6072–6079.

Miller GA (1995) WordNet: a lexical database for English. Communications of the ACM 38:39–41.

Mullally SL, Maguire EA (2011) A new role for the parahippocampal cortex in representing space. The Journal of Neuroscience 31:7441–7449.

Nagy K, Greenlee MW, Kovács G (2011) Sensory competition in the face processing areas of the human brain. PLOS ONE 6:e24450.

Nishimoto S, Vu AT, Naselaris T, Benjamini Y, Yu B, Gallant JL (2011) Reconstructing visual experiences from brain activity evoked by natural movies. Current Biology 21:1641–1646.

Op de Beeck HP, Haushofer J, Kanwisher NG (2008) Interpreting fMRI data: Maps, modules and dimensions. Nature Reviews Neuroscience 9:123–135.

Preston TJ, Guo F, Das K, Giesbrecht B, Eckstein MP (2013) Neural Representations of Contextual Guidance in Visual Search of Real-World Scenes. The Journal of Neuroscience 33:7846–7855.

Ptak R (2012) The Frontoparietal Attention Network of the Human Brain. The Neuroscientist 18:502–515.

Reddy L, Kanwisher NG (2007) Category selectivity in the ventral visual pathway confers robustness to clutter and diverted attention. Current Biology 17:2067–2072.

Reddy L, Kanwisher NG, VanRullen R (2009) Attention and biased competition in multi-voxel object representations. Proceedings of the National Academy of Sciences 106:21447–21452.

Reuter M, Schmansky NJ, Rosas HD, Fischl B (2012) Within-subject template estimation for unbiased longitudinal image analysis. NeuroImage 61:1402–1418.

Reynolds JH, Chelazzi L (2004) Attentional modulation of visual processing. Annual Review of Neuroscience 27:611–647.

Sakai K, Rowe JB, Passingham RE (2002) Active maintenance in prefrontal area 46 creates distractor-resistant memory. Nature Neuroscience 5:479–484.

Smith SM (2002) Fast robust automated brain extraction. Human Brain Mapping 17:143–155.

Verstynen TD, Deshpande V (2011) Using pulse oximetry to account for high and low frequency physiological artifacts in the BOLD signal. NeuroImage 55:1633–1644.

Wolfe JM (2012) Saved by a log: How do humans perform hybrid visual and memory search? Psychological Science 23:698–703.

